# Hybrid systems approach to modeling stochastic dynamics of cell size

**DOI:** 10.1101/044131

**Authors:** Cesar Augusto Vargas-Garcia, Abhyudai Singh

## Abstract

A ubiquitous feature of all living cells is their growth over time followed by division into two daughter cells. How a population of genetically identical cells maintains size homeostasis, i.e., a narrow distribution of cell size, is an intriguing fundamental problem. We model size using a stochastic hybrid system, where a cell grows exponentially over time and probabilistic division events are triggered at discrete time intervals. Moreover, whenever these events occur, size is randomly partitioned among daughter cells. We first consider a scenario, where a timer (i.e., cell-cycle clock) that measures the time since the last division event regulates cellular growth and the rate of cell division. Analysis reveals that such a timer-driven system cannot achieve size homeostasis, in the sense that, the cell-to-cell size variation grows unboundedly with time. To explore biologically meaningful mechanisms for controlling size we consider three different classes of models: i) a size-dependent growth rate and timer-dependent division rate; ii) a constant growth rate and size-dependent division rate and iii) a constant growth rate and division rate that depends both on the cell size and timer. We show that each of these strategies can potentially achieve bounded intercellular size variation, and derive closed-form expressions for this variation in terms of underlying model parameters. Finally, we discuss how different organisms have adopted the above strategies for maintaining cell size homeostasis.

## I. INTRODUCTION

Stochastic hybrid systems (SHS) constitute an important mathematical modeling framework that combines continuous dynamics with discrete stochastic events. SHS are increasingly being used to study noise and uncertainty not only in engineering systems [1]–[10], but also in life science applications [11]–[18]. Here we use SHS to model a universal feature of all living cells: growth in cell size (volume) over time and division into two viable progenies (daughters). A key question is how cells regulate their growth and timing of division to ensure that they do not get abnormally large (or small). This problem has ben referred to literature as *size homeostasis* [19]–[32], and is a vigorous area of current experimental research in organisms ranging from bacteria [28], [32], algae [33], [34], yeast [35] and mammalian cells [36], [37]. We investigate if phenomenological models of cell size dynamics based on SHS can provide insights into the control mechanisms needed for size homeostasis.

The proposed model consists of two state variables: *V*(*t*), the size of an individual cell at time *t*, and a timer *τ* that measures the time elapsed from when the cell was born (i.e., last cell division event). This timer can be biologically interpreted as an internal clock that regulates cell-cycle processes. Time evolution of these variables is governed by the ordinary differential equations

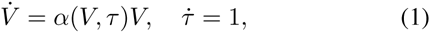

where *α*(*V*, *τ*) denotes the *growth rate* that can potentially depend on both the cell size and timer. A constant *α* implies exponential growth over time. Cell division events occur at discrete time intervals with *division rate f*(*V*, *τ*). In the stochastic formulation, the probability that the cell will divide in the next infinitesimal time interval (*t*, *t* + *dt*) is given by *f*(*V*, *τ*)*dt*. Moreover, whenever a division event is triggered, the timer is reset to zero and the size is reduced to *βV*, where *β* ∈ (0,1) is a random variable following a beta distribution. Assuming symmetric division, *β* is on average half, and its variance quantifies the error in partitioning of volume between daughters. The overall SHS model is illustrate in Fig. 1 and incorporates two important noise sources: randomness in timing of division and partitioning.

Depending on the functional forms of *f* and *α*, we consider four different strategies to control size:

1) *Timer controlled system*: Both the growth and division rates are functions of *τ* only.
2) *Size dependent growth:* Growth rate *α*(*V*) is a function of size and the division rate *f*(*τ*) is timer controlled.
3) *Size dependent division*: Constant growth rate and division rate *f*(*V*) is a function of size.
4) *Size and timer dependent division:* Constant growth rate and division rate *f*(*V*, *τ*) that depends on both variables.

**Fig. 1.**
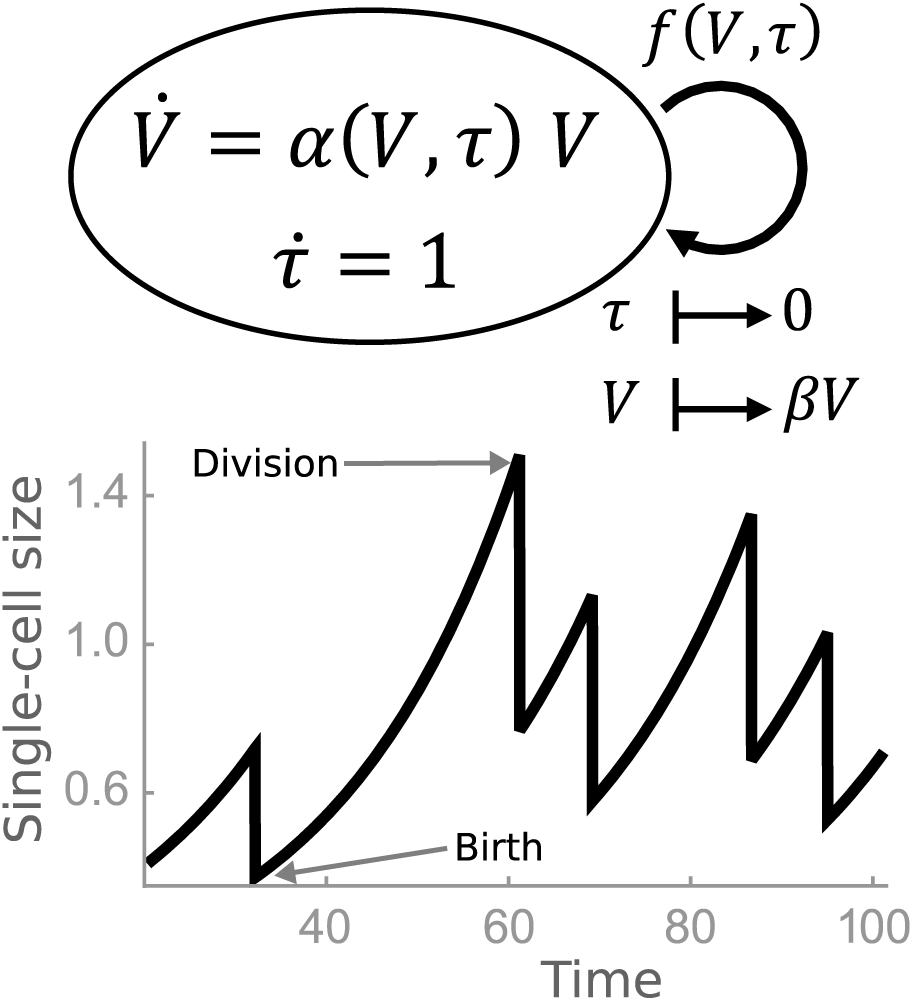
SHS model for capturing time evolution of cell size. The size of single cell *V*(*t*) grows exponentially with growth rate *α*(*V*,*r*), where r represents a cell-cycle clock that measures the time since the last division event. Cell division occurs at rate *f*(*V*,*r*), which resets r to zero, and divides the size into half. Partitioning errors in size are quantified using the random variable *β*. A sample trajectory of *V*(*t*) is shown with cycles of growth and division.

Our key contribution is to show that the first strategy of a timer controlled growth and division is not physiologically meaningful, in the sense that, the variance in cell size becomes arbitrarily large for a given mean cell size. Hence, size-controlled growth and/or division is a key requirement of size homeostasis. Analysis reveals that remaining strategies (i.e., 2 – 4 above) can result in bounded cell size variance, and we study how this variance is impacted by different noise sources. In some cases, exact analytical formulas for the first and second order moments of the cell size are derived in terms of model parameters. When exact solutions are not available, approximation moment closure methods are used to investigate cell size statistics. Finally, we discuss how diverse organisms have adopted these different strategies (or a combination of them) to achieve size homeostasis.

## II. TIMER DEPENDENT GROWTH AND DIVISION

We begin by considering a scenario, where both the growth and division rates are arbitrary functions of *τ*, but do not depend on *V*. Note that a constant growth/division rate would be a special case of this class of system. The SHS can be compactly written as

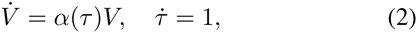

with reset maps

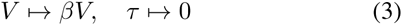

that are activated at the time of division. The probability that a cell division event will occur in the next infinitesimal time interval (*t*,*t* + *dt*) is given by *f*(*τ*)*dt*, where f can be interpreted as a “hazard function” [38]. Let *T*_1_, *T*_2_, … denote independent and identically distributed (i.i.d.) random variables that represent the time interval between two successive division events. Then, based on the above formulation, the probability density function for *T*_*i*_ is given by

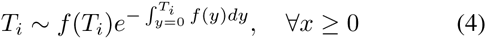

with mean duration 〈*T*_*i*_〉, where the symbol 〈 〉 is used to denote the expected value of a random variable. Note that a constant division rate in (4) would lead to an exponentially distributed *T*_*i*_. For this class of models, the steady-state statistics of V is given by the following theorem.

### Theorem 1

Consider a newborn cell with size V_0_ born at time t = 0, and goes through cycles of growth and division as per the SHS (2)-(3). Then

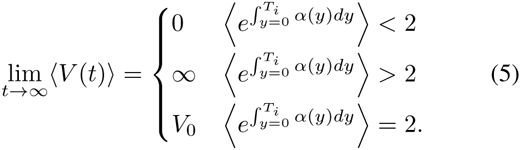

*Moreover*,

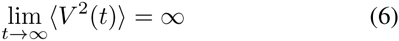

when 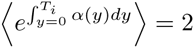.

### Proof of Theorem 1

Let *V*_*i*–1_ denote the cell size just at the start of the *i*^*th*^ cell cycle. Using (2), the size at the time of division in the *i*^*th*^ cell cycle is given by

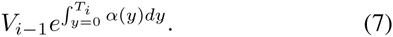

Thus, the size of the newborn cell in the next cycle is

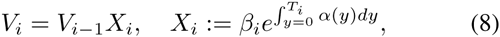

where *β*_*i*_ ∈ (0, 1) are i.i.d random variables following a beta distribution and *X*_*i*_ are i.i.d. random variables that are a function of *β*_*i*_ and *T*_*i*_. From (8), the mean cell size at the start of *i*^*th*^ cell cycle is given by

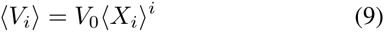

and will grow unboundedly over time if 〈*X*_*i*_〉 > 1, or go to zero if 〈*X*_*i*_〉 < 1. Using the fact that 〈*β*_*i*_〉 =0.5 (symmetric division of a mother cell into daughter cells), *β_i_* and *T*_*i*_ are independent, (5) is a straightforward consequence of (9). It also follows from (8) that

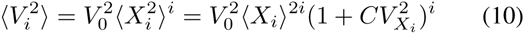

where 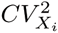 represents the coefficient of variation squared of 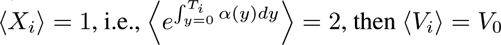 and

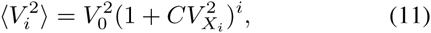

which results in (6).

In summary, above results show the unless functions *α*(*τ*) and *f*(*τ*) are chosen such that

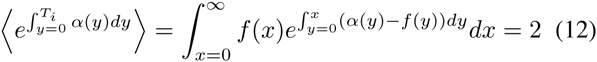

cell size would either grow unboundedly or go extinct. Moreover, in the case of a stable mean cell size, the statistical fluctuations in size would grow unboundedly. *Hence, any regulation of growth and division rates using an internal cell-cycle clock will not lead to size homeostasis*.

## III. SIZE-DEPENDENT GROWTH AND TIMER-DEPENDENT DIVISION

Recent work measuring sizes of single mammalian cells from their birth to division have found lowering of growth rates as cells become bigger [36], [37], [39]. These experimental studies suggest that size homeostasis in mammalian system relies on size-dependent regulation of growth rate. We explore what forms of such regulation will lead to bounded moments of *V*. Consider the following model

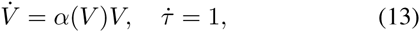

where growth rate is now size dependent and is a monotonically decreasing function with

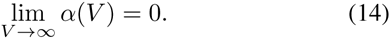

We assume that *V̇* in (13) is bounded, i.e.

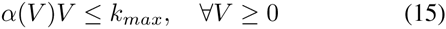

As before, division events occur with rate *f*(*τ*) and inter-division times are given by *T*_*i*_ in (4). We further consider biologically meaningful functions *f*(*τ*) that are monotonically increasing, i.e., probability of division increases as more time elapses since cell birth. The result below shows that if the growth rate for small cell size is large enough, then all moments of *V* remain bounded and converge to non-zero values.

### Theorem 2

Let growth rate *α*(*V*) satisfy (14)-(15) and *α*_0_ := lim_*V*→0_*α*(*V*) be chosen such that

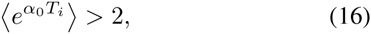

then

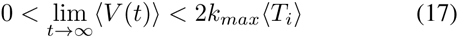

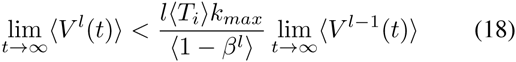

where *l* ∈ {2,3,…}, 〈*T*_*i*_〉 is the mean cell-cycle duration, and *β* ∈ (0,1) is a random variable quantifying the error in partitioning of volume between daughters:

### Proof of Theorem 2

Consider a newborn cell with sufficiently small size born at time *t* = 0. Then, the mean cell size will grow in successive generation iff the second inequality in (5) is true for *α*_0_, which results in (16). In essence, (16) ensures that cell size will not go extinct. Next we show that the mean cell size is bounded from above.

Based on Dynkin’s formula, the time evolution of statistical moments for the SHS (13) and (3) is given by

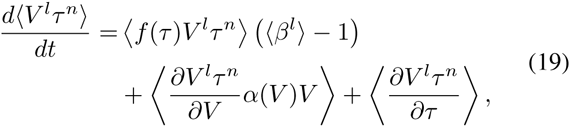

for non-negative integers *l* and *n* [40]. The dynamics of the mean cell size is obtained by setting *l* = 1 and *n* = 0

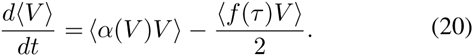

Using (15), the fact that *τ* and *V* are positively correlated and *f* is a increasing function, the above equation reduces to the following inequality

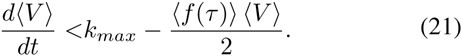

Since at steady state

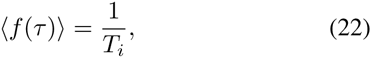

[41], (21) implies (17). From (19), the time evolution of the *l*_*th*_ order moment is obtained as

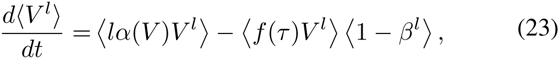

which as before, using (15) can be reduced to the inequality

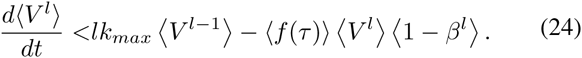

Using (22), the above inequality implies (34).

An extreme example of size-dependent growth is when *α* becomes inversely proportional to size. In this case *V* in (13) is constant, and corresponds to cells growing linearly in size as, as experimentally reported for some organisms [37], [42]. For this case, the result below provides exact closed-form expressions for the first and second-order statistical moments of *V*.

### Theorem 3

Let the growth rate be given as

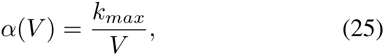

and corresponds to the following SHS continuous dynamics

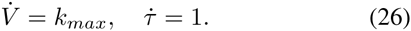

Then, the steady-state mean and coefficient of variation squared of *V*(*t*) is given by

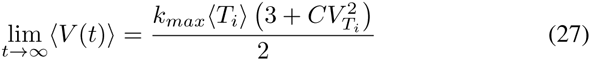

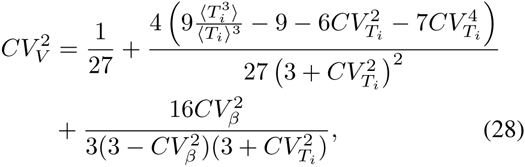

where 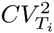. and 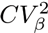 denote randomness in the interdivision times (*T*_*i*_) and partitioning errors (β), respectively, as quantified by their coefficient of variation squared.

### Proof of Theorem 3

Due to space limitations we omit the proof of this Theorem. We refer interested readers to [43], [44] where we analyzed a similar systems. More specifically, in [43] we considered linear accumulation of protein molecules inside a cell, and cell division events occurs at times *T*_*i*_ that follow a general class of distributions. Moreover, whenever the event occurs, proteins are partitioned among daughter cells based on a binomial distribution. The key difference between this theorem and [43] lies in the stochastic reset that is activated at the time of division: here, volume is reset via (3), while in [43] the state variable (protein copy number) is reset based on a binomial distribution.

Interestingly, that the mean cell size in (27) not only depends on the mean inter-division times 〈*T*_*i*_〉, but also on its second-order moment 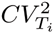. Thus, making the cell division times more random (i.e., increasing 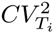) will also lead to larger cells on average. Moreover, (28) shows that the magnitude of fluctuations in cell size 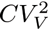 depend on *T*_*i*_ through its moments up to order three. Note that if 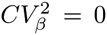 (no partitioning errors) and inter-division times approach their deterministic limit (i.e., *T*_*i*_ approaches a delta distribution *T*_*i*_ = 〈*T*_*i*_〉 with probability one), then

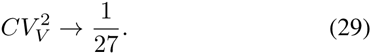

This non-zero value for 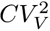 in the limit of vanishing noise sources represent variability in size from cells being in different stages of the deterministic cell cycle (i.e., some cells have just been born while others are close to division). Our result also allows for decomposing 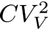 into biologically meaningful terms representing contributions from different noise sources. The terms from left to right in the righthand side of, terms from left to right in (28) represent contributions to 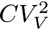 from i) Deterministic cell-cycle and ii) Random timing of division events and iii) Partitioning errors at the time of division. If fluctuations in *T*_*i*_ around 〈*T*_*i*_〉 are small, then using Taylor series

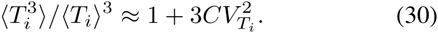

Substituting (30) in (28) and ignoring 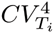 *T*_*i*_ and higher order terms yields

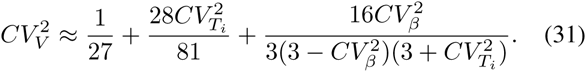

It is important to point out that since *β* follows a beta distribution with mean 〈*β*〉 = 1/2, 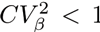 in (28) and (31). This constraint implies that 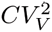 always decreases with increasing 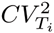, even though the last term in (31) is inversely related to 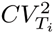.

In summary, our result show that appropriate regulation of growth rate by size (as seen in mammalian cells) can be an effective mechanism for achieving size homeostasis. We next consider a different class of models where size-based regulation is at the level division rather than growth.

## VI. CONSTANT GROWTH RATE AND SIZE-DEPENDENT DIVISION

Single cell observations in many different bacterial species indicate exponential growth (i.e., constant α) in between diviwhich implies (34) and ensures boundedness of all moments. sion events [45]–[47]. For such organisms, a size-dependent division rate is essential for maintaining size homeostasis. Towards that end, consider the following SHS continuous dynamics with constant growth rate

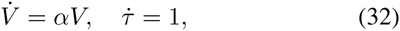

with the probability that a cell division event will occur in the next infinitesimal time interval (*t*,*t* + *dt*) given by *f*(*V*)*dt*, where f is now a monotonically increasing function of *V*. The theorem below give sufficient conditions on f for size homeostasis.

### Theorem 4

Let there exist constants k > 0 and p > 0 such that

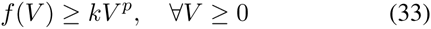

and f_0_ := lim_V→0_ f(V) < 2α, then for the SHS given by (32) and (3)

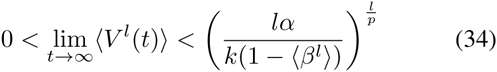

where *l* ∈ {1,2,…}.

### Proof of Theorem 4

Consider a newborn cell with *sufficiently small* size at time *t* = 0. Then, its division rate is a constant *f*_0_ and the time to division is exponentially distributed with mean 1/*f*_0_. Based on Theorem 1, the mean size will grow over successive generations (and not go extinct) for a constant growth rate iff

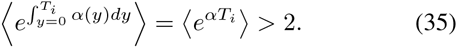

Using an exponentially distributed T_i_ with mean 1/f_0_ in the above inequality yields *f*_0_ < 2*α*, which is the necessary and sufficient condition for preventing cell size to go extinct. The time evolution of moments is given by

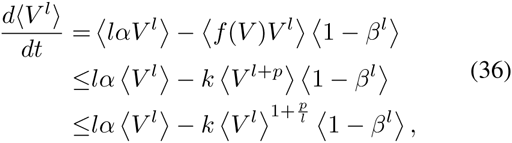

which implies (34) and ensures boundedness of all moments.

### A. The Sizer strategy

Next we analyze in detail a common example of size-dependent division, that has been referred to in the literature as the “sizer strategy” [48]–[50]. In this case, cells sense how big they are and divide when they reach a critical size threshold. Such as strategy can be implemented by choosing

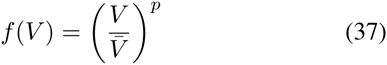

where *V̅* is a positive constant and *p* is a positive integer. A large enough *p* corresponds to division events occurring when cell size reaches a critical threshold *V̄*. Below, we show how closure schemes can be used to get approximate formals
for the moments.

Using Dynkin’s formula for the SHS defined by (32) and (37) results in the following moment dynamics

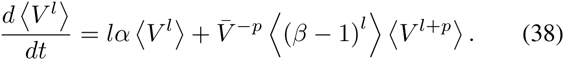

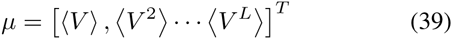

be a vector of moments up to order *L*. We refer to *L* as the order of truncation for the moment dynamics. Using (38), the time evolution of *μ* can be compactly written as

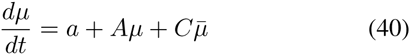

for some vector *a*, matrices *A* and *C* that are dependent on model parameters, and

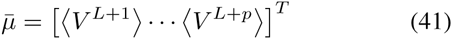

is the vector of higher order moments. Note that nonlinearities in the stochastic systems lead to the well known problem of closure, where time evolution of lower order moments μ depends on higher-order moments *μ̄*. To solve (40), we use moment closure techniques that express

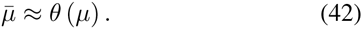

### B. Moment closure approximation

Using the closure (42), yields the following approximate moment dynamics

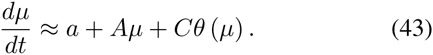

While various techniques are available to construct functions *θ* [51]–[54], we use recently proposed derivative-matching methods to close moment equations [17], [55]. For example, consider *L* = 2 in (39) (second order of truncation) and *p* = 2, then the higher-order moments are given by

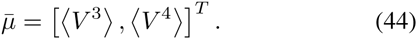

Based on derivative-matching, these higher-order moments can be approximated in terms of lower-order moments as

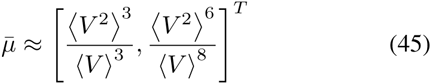

[17], [55]. It turns out that the closed moment equations (43) corresponding to *L* = 2 and derivative-matching closure yield analytical expressions for the steady-state moments for any integer *p* in (37). In particular, the steady-state mean and coefficient of variation squared of cell size is given by

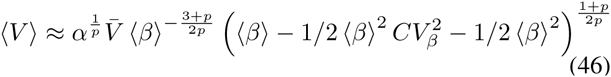

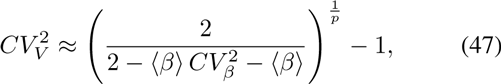

respectively. For symmetric division (〈*β*〉 = 1/2), above equations reduce to

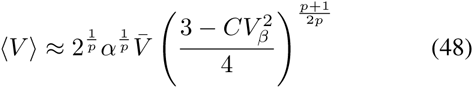

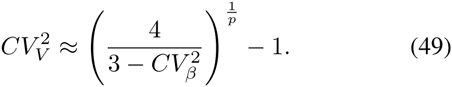

These results qualitatively reproduce the behavior (as *p* increase, noise in volume decrease) seen from computationally expensive Monte Carlo simulations of the SHS (Fig. 2).

Intriguingly, (48)-(49) show that the mean cell size is affected by 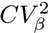 (magnitude of error in partitioning of volume between daughter cells): larger partitioning errors decrease the average cell size, but increase 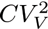 (see Fig. 3). Another novel insight from (49) is that 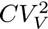 is independent of cell growth rate (*α*) and cell size threshold *V̅*. Indeed, computing 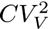 from Monte Carlo simulation results in the exact same noise curve as in Fig. 2 for different values of *α* and *V̄*.

**Fig. 2.**
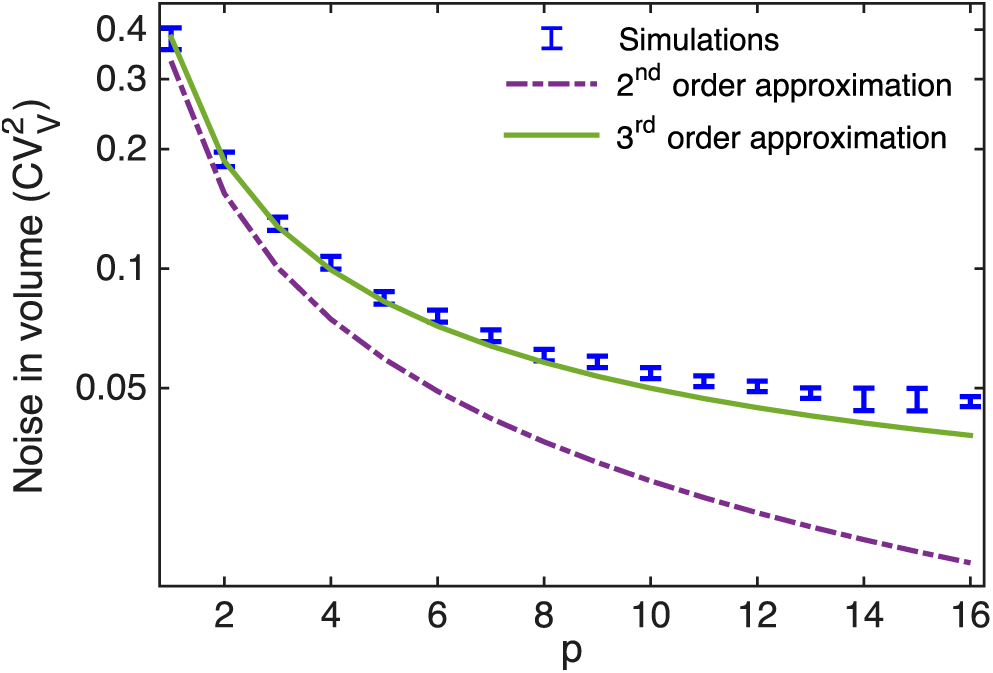
Noise in the cell size (measured by the coefficient of variation squared) as a function of the parameter *p* for division rate given by (37). Comparisons are shown between results obtained via Monte Carlo simulations and using closed moment equation (43) corresponding to 2^*nd*^ and 3^*rd*^ order truncation. While the qualitative behavior is similar, increasing the order of truncation leads to a better quantitative match to results obtained from Monte Carlo simulations. Partitioning noise is assumed to be zero.

**Fig. 3.**
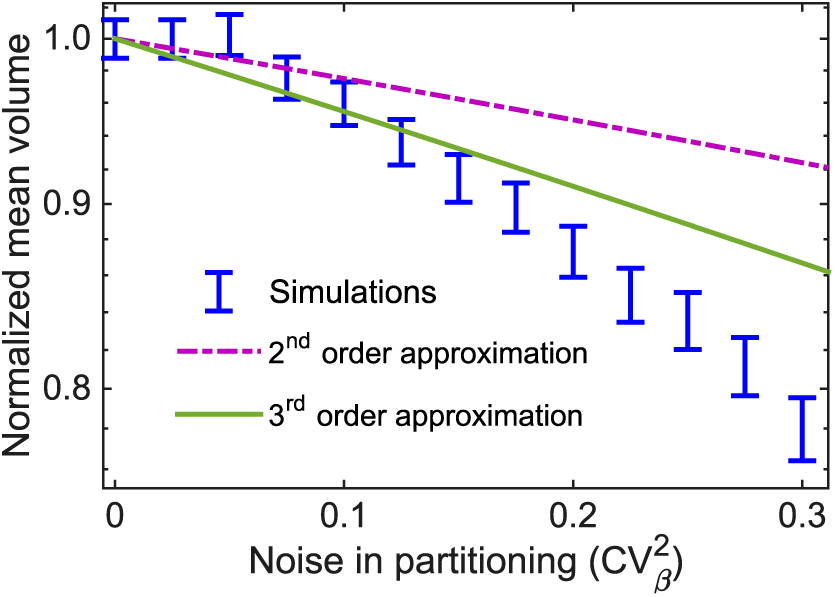
Mean cell size decreases with increasing noise in partitioning (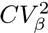). Average size obtained by running a large number of Monte Carlo simulations is shown together with corresponding solutions obtained from closed moment equations for 2^*nd*^ and 3^*rd*^ order truncation. Mean cell size was normalized by the average size in the case of zero noise in partitioning (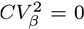). Parameters used were *α* = 0.05 min^‒1^, *V̅* = 2.5 *μm*, and *p* = 2.

## V. CONSTANT GROWTH RATE, SIZE AND TIMER DEPENDENT DIVISION

In this section, we consider a constant growth rate (as in the previous section) but with division rates *f*(*V*, *τ*) that take information from both size and cell-cycle clock. Before providing an examples of this form of regulation, we briefly discuss sufficient condition on *f*(*V*, *τ*) for size homeostasis.

For a sufficient small cell size, division rate is given by *f*_0_(*τ*) := lim_*V*→0_ *f*(*τ*, *V*). Then from Theorem 1, a necessary and sufficient condition for cell size to not go extinct is

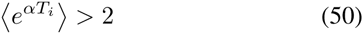

where the distribution of *T*_*i*_ is given by (4) with f replaced by *f*_0_. Moreover, a sufficient condition that ensures moments of *V* remain bounded is

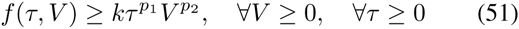

for some positive constants *k*, *p*_1_ and *p*_2_. Given the space constraints we omit the proof of this result. Next, we analyze in detail a recently uncovered size homeostasis strategy, where the division rate is a function of both size and cell-cycle clock.

### A. The Adder strategy

Recent observations is many bacterial species have found that timing of cell divisions are regulated so as to add a fixed size from cell birth to division [21], [25], [27], [28], [56]–[59]. Since cell size grows exponentially (i.e., constant α), at any given time t the size added since birth would be given by

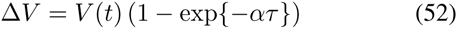

which is determined by both *V* and *τ*. As per this “adder strateg”, division is triggered when Δ*V* reaches a threshold. Below, we discuss how this “adder strategy” is implemented and perform a stochastic analysis of the model using closure techniques.

A molecular implementation of the adder model was proposed by [26], [57], [60] and shown in Fig. 4. It consists of a time-keeper protein with level M that is produced at a rate proportional to size. The continuous dynamics is now given by

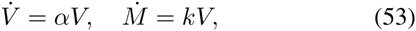

where *k* is a positive proportionality constant. Cell division event occurs when *M*(*t*) reaches a threshold *M*, and whenever the even occurs state variables are reset as

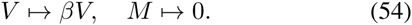

Based on this formulation the volume added between two division events will be fixed and given by

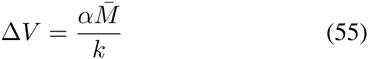

[26]. To practically implement the “adder strategy” we consider the SHS (53)-(54) with a division rate

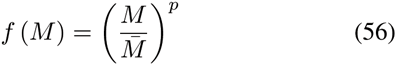

for a positive integer *p*, where large enough *p* would correspond to division occurring when *M*(*t*) = *M̅*. We next explore the moment dynamics of this systems using closure schemes.

**Fig. 4.**
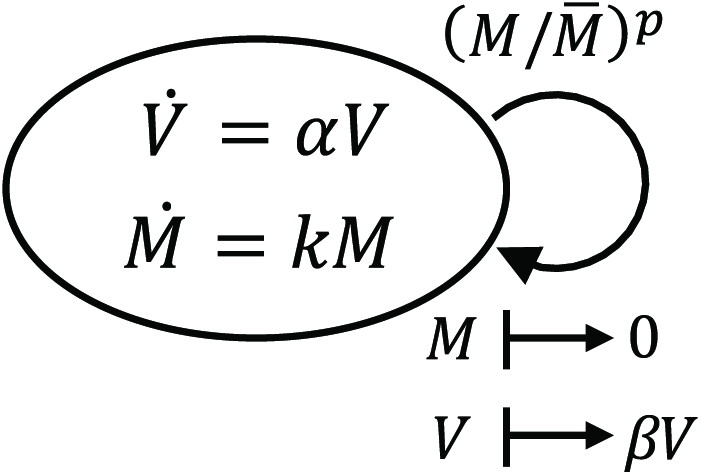
SHS representation of a molecular implementation of the adder strategy. A time-keeper protein (*M*) is produced at a rate proportion to size, an division is triggered when *M* reached a threshold *M̅*. Just after division, the protein is fully degraded. Based on the implementation, the size added between two successive events is constant and given by (52).

**Fig. 5.**
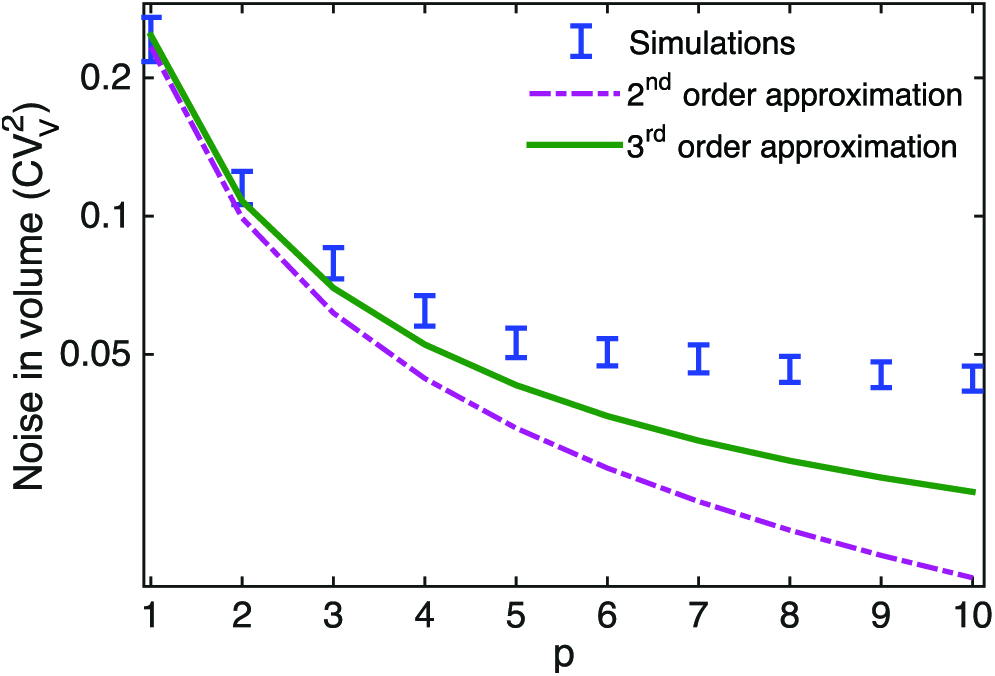
Noise in the cell size (measured by the coefficient of variation squared) as a function of the parameter *p* in Fig. 3. 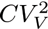 decreases with increasing *p* as illustrated by Monte Carlo simulations and solutions of closed moment equations based on a 2^*nd*^ and 3^*rd*^ order truncation. Increasing the order of truncation results in a better match with Monte Carlo simulations.

### B. Moment closure approximation

For the SHS (53)-(54), the time evolution of the statistical moments is obtained as

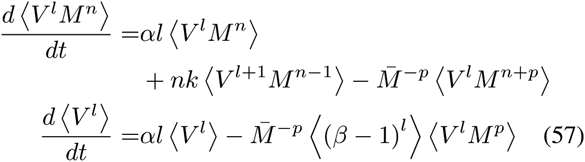

for *l*, *n* ∈ {0,1, 2,…,}. The moment equations can be closed using derivative-matching technique, which expresses any higher-order moment as

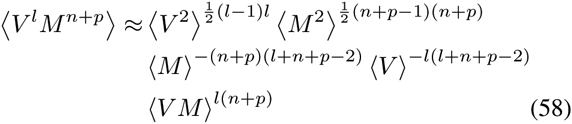

in terms of moments of order up to two [55]. Closing the moment dynamics of first and second order moments (i.e., second order truncations) yields the following steady-state mean and noise levels for cell size

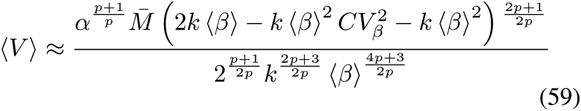

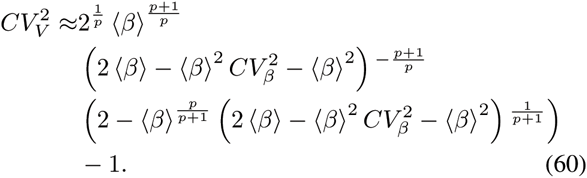

In the case of symmetric division (〈β〉 = 1/2), these equations reduces to

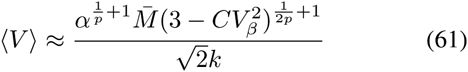

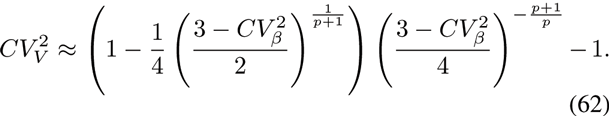

Above results show that as observed earlier, increasing partitioning errors will lead to smaller mean cell size but larger noise. Moreover, 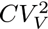 monotonically decreasing with increasing *p* consistent with results from Monte Carlo simulation (Fig. 5). A comparison of Fig. 3 and 5 also reveal that for a given value of *p*, the adder provides lesser noise in size compared to the sizer. Finally, equation (60) shows that 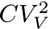 is independent of cell growth rate (*α*), and parameters *k* and *M̅*.

## VI. Conclusion

Here we have used a SHS framework to model time evolution of size of a single cell undergoing cycles of growth and division. Our goal was to uncover mechanisms responsible for size homeostasis, i.e., ensuring a tight distribution of cell size. The model is defined by three features: a growth rate *α*(ν, *τ*), a division rate *f*(*V*, *τ*), and a random variable *β* ∈ (0,1) that determines the reduction in size when division occurs. Results show that growth/division rates that only depend on the cell-cycle clock *τ* are not physiologically, since they lead to unboundedly increasing cell size variance (Theorem 1). The key contribution of this work is to identify sufficient conditions on growth/division rates that prevent extinction of cell size (i.e., 〈*V*〉 → 0 as *t* → 0) and also lead to bounded cell size variance (Theorems 2-4).

We also analyzed two commonly used models for size homeostasis: the sizer (cell division occurs at a critical size) and the adder (cell division occurs after adding a critical size from cell birth). Interestingly, closure techniques were shown to yield approximate analytical formulas for the first and second order moments of cell size. There formulas reveal a novel result that increasing degree of partitioning errors 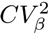 in these models leads to smaller cells on average and higher noise in cell size 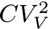. Moreover, for many parameter regimes the adder strategy was found to result in a lower 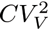 than the sizer.

In summary, theoretical tools for SHS can provide fundamental understandings of regulatory mechanisms maintaining size homeostasis. Future work will focus on developing computational tools for inferring the growth and division rate functions from single-cell measurements of cell size over many generations. It will be interesting to apply these tools to data, and infer these functions in the context of different biological organisms.

## ACKNOWLEDGMENT

AS is supported by the National Science Foundation Grant DMS-1312926.

